# Targeted and bilateral blood flow monitoring in middle cerebral artery using diffuse correlation spectroscopy

**DOI:** 10.64898/2026.09.21.750817

**Authors:** Kavita Sharma, Susweta Das, Kimberly Gonsalves, Soumyajit Sarkar, U.S. Srinivasan, Hari M Varma

## Abstract

**Objective:** To develop and validate a dual-probe Diffuse Correlation Spectroscopy (DCS) system for non-invasive and simultaneous, monitoring of cerebral blood flow (CBF) in the bilateral Middle Cerebral Artery (MCA) territories, and expanding the utility of conventional DCS limited to cortical-volume-based CBF measurements to vessel-specific cerebral perfusion monitoring.

**Methods:** A dual-probe DCS system was designed for non-invasive monitoring of MCA-specific perfusion. Probe placement and protocol optimization study has been performed using anatomical landmarks, motor and speech activation tasks in healthy volunteers. System stability and repeatability were further evaluated in a pilot cohort of 30 healthy (age, 25 ± 7 years) participants using optimized probe position and protocol. A bilateral MCA ischemic Lacunar Infract stroke case report also validated the feasibility of the system in clinical settings.

**Results:** Measurements demonstrated superior sensitivity towards MCA-territory perfusion at targeted probe locations compared to off-MCA positions. In pilot cohort, significant increase of 30.34 ± 21.56% and 36.48 ± 21.22% in rCBF corresponding to hand squeeze and speech task respectively showed reproducible physiological responsiveness of the system (p *<<* 0.001). Measurement done on a patient with bilateral MCA ischemic Lacunar Infract stroke showed a significant change of 30% during speech for both the MCAs but no significant change is observed for hand squeeze tasks.

**Conclusion:** The custom built dual-probe DCS system enables non-invasive, operator-independent, targeted and continuous monitoring of rCBF within bilateral MCA territories.

**Significance:** This approach enables the potential use of DCS system for bilateral and vessel-specific monitoring of cerebral perfusion in the MCA territories.

## I. Introduction

MCA is the largest branch of the internal carotid artery (ICA) located within the lateral sulcus (Sylvian fissure) between the frontal and temporal lobes and plays a vital role in maintaining cerebral blood circulation. As a key component of the circle of Willis, it supplies oxygenated blood to the lateral aspects of the frontal, parietal, and temporal lobes, along with important deep brain structures such as the basal ganglia and internal capsule [1]. Anatomically, the MCA is divided into four segments—M1 (sphenoidal), M2 (insular), M3 (opercular), and M4 (cortical)—which give rise to cortical and deep perforating branches that support regions responsible for motor, sensory, language, and cognitive functions [2]. Due to its extensive vascular territory, the MCA is the most frequently affected artery with ischemic stroke [3].

Stroke remains one of the leading causes of mortality, long-term disability worldwide and is increasingly prevalent among younger populations [4], [5]. Rapid assessment and continuous monitoring of cerebral perfusion are critical for improving clinical outcomes, particularly during the acute phase following stroke onset [6]. Current clinical practice relies on radiological imaging modalities such as Computed Tomography (CT) and Magnetic Resonance Imaging (MRI) for initial diagnosis and treatment planning [7]. While these techniques provide high spatial resolution, they are not suitable for continuous bedside monitoring of cerebral flow dynamics, which is crucial during the first 6-48 hours post-stroke when patients are most vulnerable to secondary injury [8].

Transcranial Doppler (TCD) ultrasound is the current point-of-care modality for assessing cerebral hemodynamic in stroke patients, particularly in major anterior circulation vessels such as MCA [9]. TCD offers real-time measurements of blood flow velocity and is non-invasive; however, it suffers from several inherent limitations, such as variability in skull bone thickness of temporal region across patients, dependence on operator expertise, and restrictions imposed by probe geometry [10]. Moreover, as a hand-held technique, TCD does not permit continuous, long-term bedside monitoring, limiting its utility in dynamic clinical settings such as Intensive Care Units (ICUs) [11]. These limitations highlight the need for a portable, non-invasive, and operator-independent technology capable of providing continuous, real-time monitoring of cerebral blood flow in a vessel-specific manner.

DCS has emerged as a promising optical technique for non-invasive quantification of microvascular blood flow that utilizes near-infrared light to probe temporal fluctuations caused by moving red blood cells (RBCs), enabling continuous, tracer-free measurement of relative CBF [12]. Previous studies have demonstrated the utility of DCS in a range of applications, including functional brain activation, neonatal monitoring, traumatic brain injury, and stroke research [13], [14]. However, most existing DCS implementations rely on diffuse probe placement over generalized cortical regions such as pre frontal cortex (FP1 and FP2), sensorimotor cortex and temporal cortex limiting their ability to provide targeted measurements from specific vascular territories such as the MCA [15].

In this study, an in-house developed DCS-based system integrated with specially designed dual probe is presented for targeted and continuous monitoring of MCA-specific relative flow perfusion. A key feature of this system is its ability to perform simultaneous bilateral monitoring of the left and right MCAs, eliminating the need for probe repositioning and thereby reducing measurement variability. Tasks perturbing MCA blood supply are performed with healthy volunteers for probe position and protocol optimization study. A measurable and reproducible changes in MCA blood flow for 30 subjects have been demonstrated using the optimized probe position and protocol. Measurement done on patient with ischemic Lacunar Infract stroke validated the feasibility of the system in clinical settings.

## II. DCS SYSTEM AND DUAL PROBE

### A. In-house DCS system design

In-house DCS system was developed containing fiber-coupled coherent laser sources (S1 and S2) with wavelength 785nm for illumination and Single Photon Avalanche Diode detectors (D1 and D2) to collect the backscattered light from the respective hemispheres [16]. A custom-designed dual probe was developed that consists of two identical sections containing one source fiber (multimode fiber) and one detector fiber (single mode fiber) each as discussed in detail in section II-B.

The laser light delivered to the MCA territory, interacts with moving red blood cells (RBCs) in the cerebral vasculature and undergoes multiple scattering. These dynamic scattering events induce temporal fluctuations in the detected light intensity. The backscattered photons were collected at source-detector separation of 17 mm. The detected signals were acquired using a National Instruments data acquisition (NI-DAQ) system interfaced with a computer for real-time signal acquisition and visualization as shown in Fig. 1. Throughout the study, power of the laser source is kept well within the safety limits as per the ANSIZ136.1 laser safety standards.

**Fig. 1.**
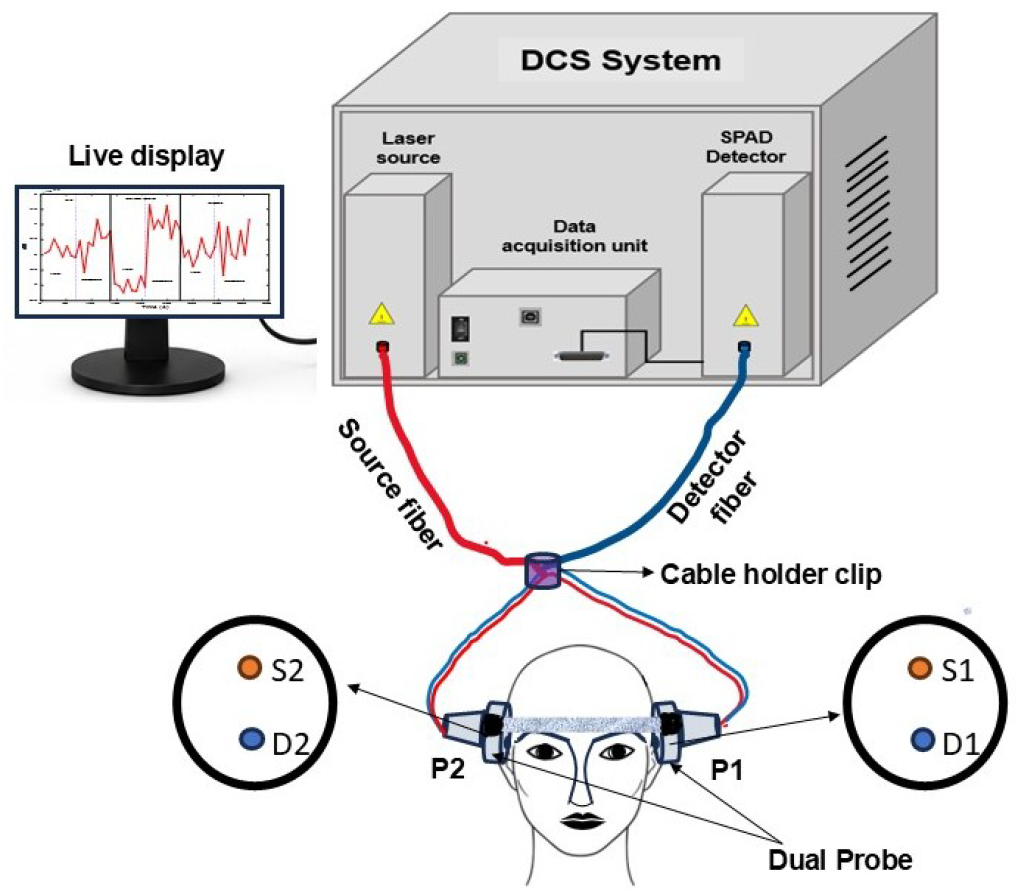
Schematic diagram showing an in-house developed DCS system integrated with dual probe (P1 and P2) for simultaneous monitoring.

### B. Custom-designed dual probe

A specific dual probe assembly is designed for simultaneous monitoring of flow dynamics in both the MCA. This dual probe assembly contains two separate, but identical probes sections denoted as P1 and P2. These probe sections can be used to monitor left MCA (LMCA) and right MCA (RMCA) respectively and vice-versa. Both the probes are connected to each other with flexible elastic bands containing sliding hooks. Since the precision in the probe placement is the key factor, the probes were designed with a transparent window to ensure the proper sampling of MCA territories.

To fabricate this transparent window, the silica-based polymer polydimethylsiloxane (PDMS) was used with curing agent (10:1). Source and detector fiber tips were embedded in the PDMS poured in 3D printed mold (white circular part as shown in Fig. 2). Subsequently, the mold containing PDMS was desiccated for 1 hr to remove all the bubbles and to get the clear vision, followed by baking at 70° C for 45 mins. Similarly, the other probe section (P2) was made and both the probes were joined with two different types of bands as shown in Fig. 2 (a). We have used a broader band (black) of width 25 mm with webbing hook. This hook ensures the stability of the probe on head and it help to easily put and remove the probe by losing and tightening the strap. The front band is kept of width 10 mm with adjustable sliding hook. A narrow strap avoids the discomfort to the eyes as it fixes over the forehead. Since, the circumference of skull varies person to person result in varying lateral distance between left and right MCA, the sliding hook gives the freedom to adjust the strap according to forehead size of the subject that helps in proper placement of probe on both MCAs simultaneously. The probe can be seen on one of the volunteer’s head from Fig. 2 (b) and (c). This probe has been designed considering its integration with ancillary support systems such as Non-Invasive Ventilation (NIV) mask that that may increase its suitability for ICU settings.

**Fig. 2.**
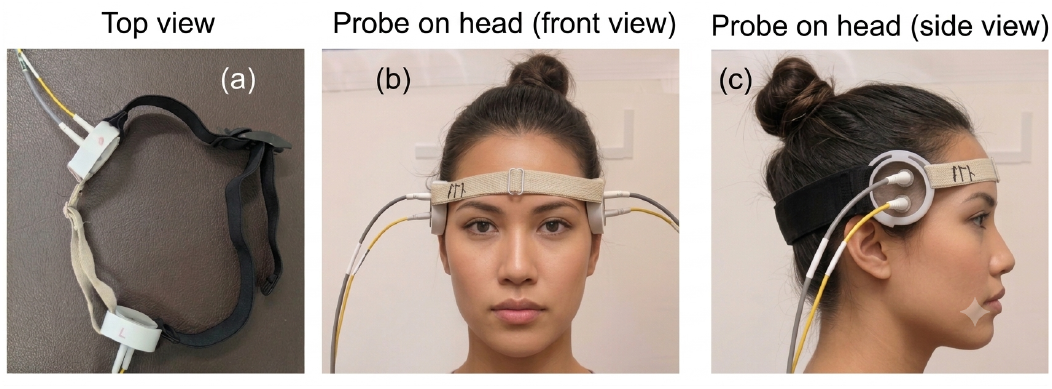
Designed and fabricated dual probe for simultaneous monitoring of both the MCA (a) top view of probe (b) front view (c) zoomed side view. (Fig. 2 (b) and (c) are AI generated images with fabricated dual probe)

## III. METHODOLOGY

## A. DCS theory

Backscattered photons detected by the SPAD were continuously recorded using a 3 sec of data acquisition window. A multi-tau autocorrelation algorithm was used to calculate the normalized intensity autocorrelation function from photon count fluctuations [17]. The electric-field autocorrelation function *g*_1_(*τ*) was obtained using the Siegert relation, *g*_2_(*τ*) = 1 + *β* | *g*_1_(*τ*) |^2^, where *β* is the coherence factor determined by the detection optics. The blood flow index (BFI), defined as *αD*_*B*_, was estimated by fitting the measured *g*_1_(*τ*) to the analytical solution of the Correlation Diffusion Equation (CDE) for a semi-infinite homogeneous medium. The CDE is given by [18],

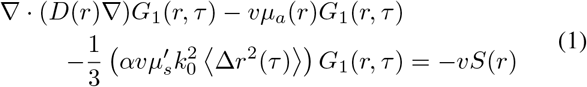

where, *v* and *τ* stands for velocity of light and delay time respectively. *D* is the photon diffusion coefficient, given as. 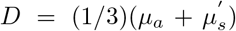 *µ*_*a*_ and 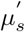 are the absorption and reduced scattering coefficients determined from an in-house time-domain NIRS system that resembles with the literature reported [10]. The RBC motion was modeled using the Brownian diffusion model, with *D*_*B*_ as the effective diffusion coefficient. The BFI *αD*_*B*_ was plotted in real time to visualize blood-flow changes during the task. Photon counting and autocorrelation were implemented in C, while acquisition and CDE fitting were performed in MATLAB R2021b.

### B. Optimization of probe position

To monitor the flow dynamics in MCA, the targeted MCA position lies within the lateral sulcus (Sylvian fissure) between the frontal and temporal lobes through which MCA traverses [19]. Pterion region (black dotted circle) including M2-M3 segment is the most appropriate region for MCA sampling (refer Fig. 3 (a)), here, M2 (Insular) ascends and travels across the surface of the insular cortex and M3 (Opercular) curves and loops over the inner margins of the frontal, parietal, and temporal opercula. As the pterion region was avoided due to hair constraint, M2 segment (black square) is considered as the targeted MCA region. In this region, relatively thin temporal bone (approximately 3–4 mm) provides close proximity to the MCA from surface, result in an optimal probe positioning ensures maximum sampling of the arterial flow with minimal interference from the surrounding cortical tissues [20]. Fig. 3 (b) shows a schematic of anatomical landmarks to approach the targeted MCA position that offers an ease in optimal probe placements. Here, T denotes the tragus, ZA is the zygomatic arch (dotted arc), G is the glabella and N is the nasion [21]. L1 is a line connecting the points T and N, L2 is a line passing through T and perpendicular to L1. L3 is a line parallel to L2 at a distance of 15 mm towards the ZA.

**Fig. 3.**
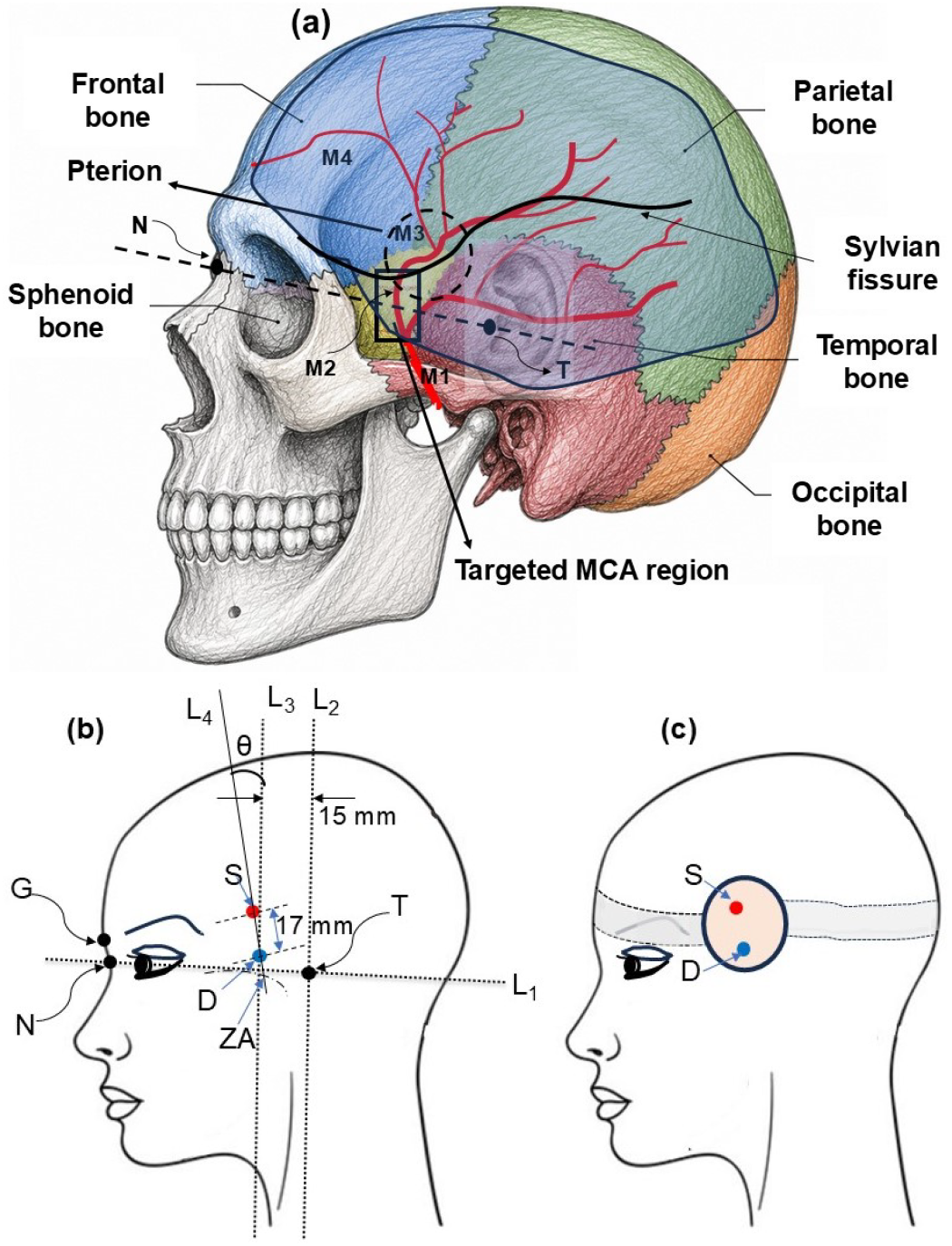
Schematic showing (a) neurocranium with various MCA branches M1-M4 along with the targeted MCA arterial region sylvian fissure i.e., M2 (black square), (b) the anatomical landmarks (sagittal view) to approach precise probe placement, and (c) the optimized DCS probe positioning for efficient sampling of flow dynamics in MCA.

The source and detector fiber tips are denoted by S (red spot) and D (blue spot) respectively. For optimal probe positioning, D should be placed on line L3 and just 5 mm above the zygomatic arch with a tolerance of ±0.5 mm and, S should be placed vertically above D such that the source detector separation is 17 ± 2.5 mm. The probe should be oriented in such a way that the line (say L4) joining both the fiber tips (source and detector) is making an angle (*θ*) of 10 degree with L3 with a tolerance of ± 2 degree. Optimized probe location/position can be seen from Fig. 3 (c). For the confirmation of MCA position, a blood flow monitoring experiment is performed at targeted MCA position along with two MCA-off positions. One position is slightly up to the targeted MCA lies at frontal bone and second probe is slightly down to the targeted MCA position that lies at zygomatic bone as shown in Fig. 4 (a). There is only lateral shift in probe position, keeping angle *θ* same for all three probe positions.

**Fig. 4.**
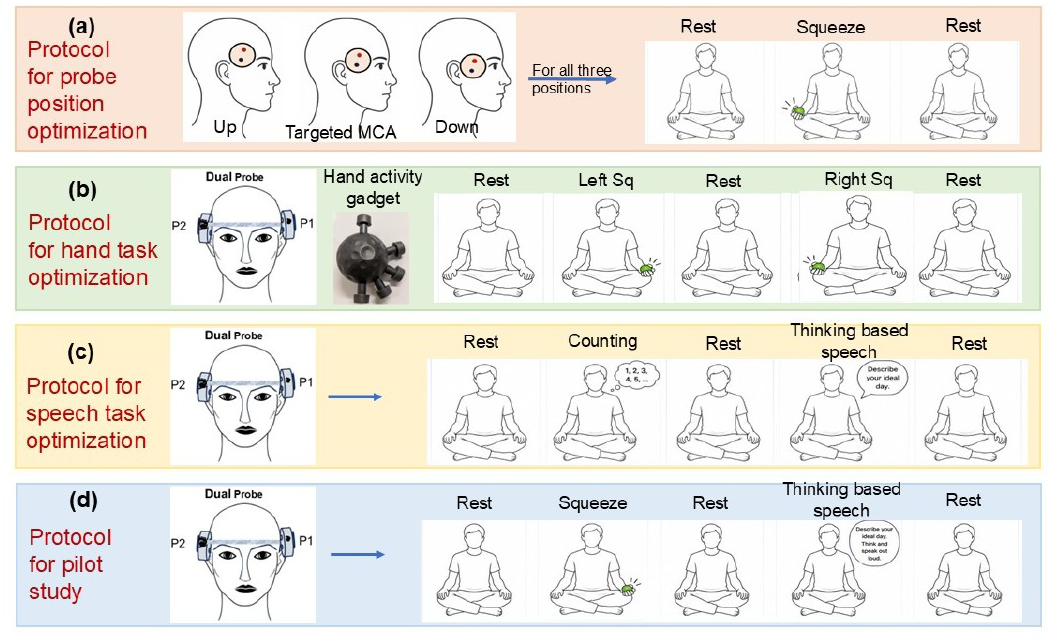
Schematic representation of (a) protocol for optimal probe position out of three selected positions: MCA targeted position, off to the MCA targeted position (b) protocol for hand task optimization (c) protocol for speech task optimization, and (d) protocol for pilot study.

### C. Protocol optimization

To carry out protocol standardization study, functional tasks that consistently perturb CBF in MCA were identified. The regions supplied by MCA include the lateral sensorimotor cortex (contains the lateral parts of the primary motor cortex, the primary somatosensory cortex, and the ventral premotor cortex) controls voluntary movements and sensations for the face, lips, jaw, and hands (motor activities), Broca’s area (expressive speech) and Wernicke’s area (receptive language comprehension) controls speech centres [22], [23].

As MCA supplies to the part of the motor cortex that controls the lower arm and hand motor activities, the fist squeezing was incorporated to perturb the flow in MCA [16]. In order to optimize the targeted MCA probe position as explained in section III-B, volunteers were asked to perform the protocol as shown in Fig. 4(a) i.e., rest (90 sec), squeeze (with contralateral hand for 60 sec), and rest (90 sec). This study was repeated with three volunteers with age group of 25 ± 3 years (2 female, 1 male). Since, left MCA and right MCA predominantly supplies to contralateral upper limb, contralateral and ipsilateral hand squeeze activity was considered for hand task optimization protocol along with a hand grip strengthener gadget as shown in Fig. 4(b).

Speech task involves contribution of both the MCAs, as right MCA is responsible for planning and left MCA is responsible for execution of speech task [17]. In order to optimize the task that shows more representation in both MCAs, two speech tasks were chosen; one of them is counting, that involves minimal thinking and another task is a question & answer task that involves thinking/planning and execution as shown in Fig. 4 (c). A total of 7 subjects (3 male and 4 female) with age (25 ± 4) years participated for the speech protocol optimization experiment. Of them, 2 data sets were excluded because the motion artifact resulted in an unreliable assessment of the changes in blood flow induced by the task.

During probe position and protocol optimization study, simultaneous monitoring of both the MCAs leads to decreased temporal resolution, approximately half to the individual MCA monitoring (tres = 8.8 sec). As the proposed dual probe is capable of monitoring blood flow individually and simultaneously both (as per the need), data is taken with one probe, keeping other probe OFF, and vice - versa. This consideration offered an ease in performing the task for the subjects and avoided the probe placement error while repositioning.

In order to check the consistency and accuracy of the system, a pilot study has been done. The protocol for pilot study involves contralateral hand squeeze activity with the hand grip strengthener gadget and the thinking-based speech task as shown in Fig. 4 (d). The experiment was performed by 30 healthy volunteers (male to female subject’s ratio 1:1) with age group of 25 ± 7, all right-handed (coincidently) and mixed ethnicity were recruited. Exclusion criteria included a history of stroke, neurodegenerative disorders, neurological conditions such as epilepsy, recurrent migraine, any musculoskeletal or neurological impairment and any hand injury affecting the upper limbs that could interfere with the experimental tasks.

### D. Consent and Ethical approval

All experimental procedures involving human participants were approved by the Institutional Ethics Committee (IEC) of the Indian Institute of Technology Bombay under protocol number IITB IEC/2024/10. The study was conducted in accordance with the principles of the Declaration of Helsinki. Written informed consent was obtained from all participants prior to their enrolment.

### E. Data analysis

For each subject, the relative Cerebral Blood Flow or rCBF (%) was quantified with respect to the baseline that is given as,

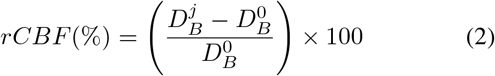

where 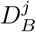 is the BFI corresponding to the j-th timestamp and 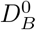 is the BFI of baseline which corresponds to the mean of all rest positions.

### F. Statistical analysis

All statistical analyses were performed using MATLAB R2021b (MathWorks, USA). Prior to hypothesis testing, the distribution of the mean rCBF values was assessed using the Lilliefors test for normality check. Since, the data satisfies the assumption of normality, significance between base to task and across different tasks (hand task and speech task) was performed using the paired t-test. Statistical significance was defined as the p value to be less than 0.01. All reported p-values are two-sided. Power analysis was performed with G*Power 3.1 software to estimate the minimum sample size required. The analysis assumed a significance level of *α* = 0.01, a statistical power (1 *− β*) of 0.95, and an expected effect size (d), derived from protocol optimization data (mean ± Standard Deviation rCBF %).

Based on these assumptions, the estimated minimum sample size considered is 10, as given in Table I. Nevertheless, 30 healthy participants were ultimately recruited to further enhance the statistical power and reliability of the study findings.

**Table I.** Power analysis for estimation of minimum sample size requirements.

| Position | Task | rCBF Mean % | rCBF SD % | d = Mean/SD | Sample size |
| --- | --- | --- | --- | --- | --- |
| Left MCA | Squeeze | 137.31 | 71.67 | 1.92 | 9 |
|  | Speech | 72.13 | 35.58 | 2.03 | 8 |
| Right MCA | Squeeze | 110.34 | 54.72 | 2.02 | 8 |
|  | Speech | 103.11 | 58.74 | 1.76 | 10 |

## IV. Results

### A. Observations of optimal probe localization study

For the validation of accurate probe localization study, the experiment was performed with three healthy human volunteers for left and right MCA, following the anatomical landmarks and protocol as discussed in the section III-B and III-C respectively.

The obtained data can be seen from rCBF plots and mean rCBF bar graphs from Fig. 5(a) and (b), for LMCA and RMCA respectively. For targeted MCA position, an increase in rCBF of 92.17 ± 24.38% and 44.07 ± 18.38% is measured during squeeze task for LMCA and RMCA respectively as compared to rCBF change for Up position (−8.16 ± 4.62% and −1.13 ± 8.87%) and Down position (6.64 ± 9.37% and 5.21 ± 8.30%). A comparative analysis revealed significantly visible increase in rCBF for targeted MCA position i.e., sylvian fissure. Consequently, targeted MCA position is considered as optimal probe position for further measurements.

**Fig. 5.**
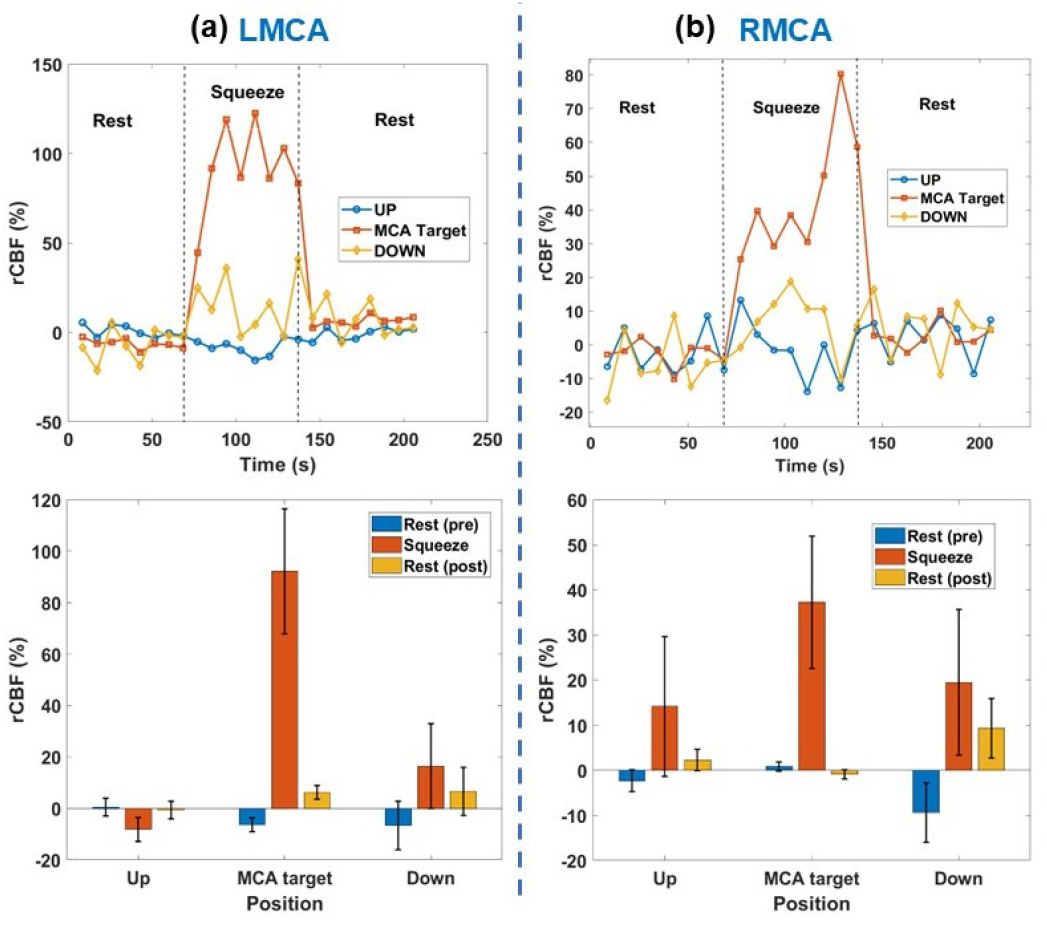
Live plot and bar graphs showing the rCBF change corresponding to contralateral hand squeeze for three different positions of DCS probe in (a) Left MCA (b) Right MCA.

### B. Observations of protocol optimization study

Protocol optimization study was performed with the optimized probe position, as discussed in section III. C, by following the protocols shown in Fig. 4 (b) and (c) for squeeze task and speech task optimization respectively. Fig. 6 (a) represents rCBF change corresponding to contralateral and ipsilateral hand squeeze task for left and right MCA. Shading part along with the plot is showing the variance. For LMCA, measured values of mean rCBF are 55.01 ± 24.07% for left squeeze (ipsilateral) and 137.31 ± 71.67% for right squeeze (Contralateral). Similarly, for RMCA, 110.34 ± 54.72% and 36.14 ± 11.19% mean rCBF is obtained for left squeeze and right squeeze respectively. Comparative analysis showed significantly higher response for contralateral hand squeeze activity in MCA territory and therefore validating it as optimal hand activity for pilot study.

**Fig. 6.**
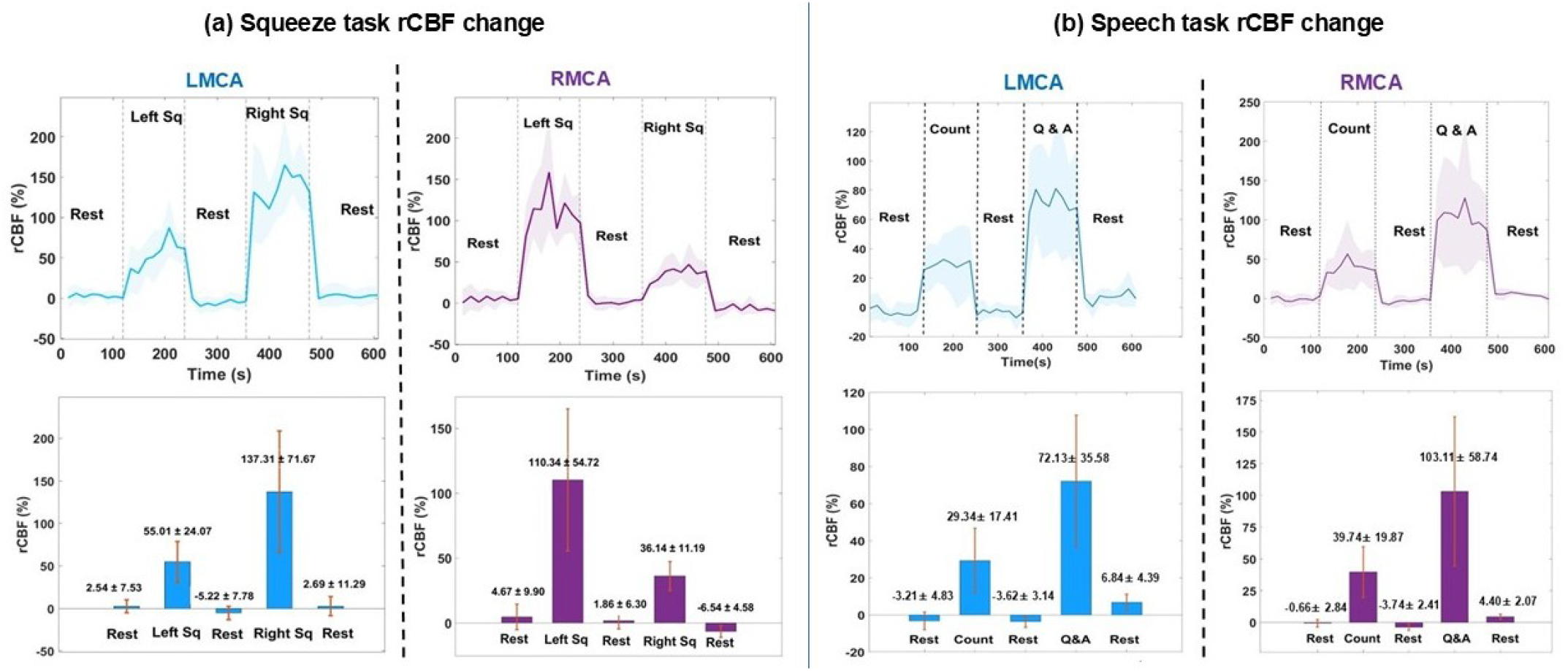
The rCBF change for (a) contralateral and ipsilateral hand squeeze task, and (b) shows rCBF change for counting and Q&A speech tasks for left and right MCA corresponding to five healthy subjects (n=5).

Furthermore, for speech task optimization study as shown Fig. 6 (b), mean rCBF obtained is 29.34 ± 17.41% for counting, 72.13 ± 35.58% for Q&A for LMCA, and 39.73 ± 19.87%, 103.19 ± 58.74% is obtained for counting and Q&A respectively for RMCA. It can be concluded from plots, bar graphs and measured data that the rCBF change is significantly higher for thinking-based speech task i.e., Q&A corresponding to both left and right MCA. Consequently, Q&A will be considered as the speech task for pilot study.

### C. Pilot study

After finalizing all parameters, the system’s consistency and accuracy was validated with 30 healthy volunteers with optimized probe position and protocol as discussed in section III. B & C. Subjects were instructed to sit relaxed and asked to perform the protocol as shown in Fig. 4 (d).

Fig. 7 (a) and (b) present the percentage change in relative cerebral blood flow (rCBF) with respect to baseline for the LMCA and RMCA, respectively, while Fig. 7 (c) and (d) show the corresponding mean rCBF values. Measured values of mean rCBF for LMCA are observed to be 30.34 ± 21.56% and 36.48 ± 21.22% for contraleteral squeeze and Q&A tasks respectively w.r.t baseline.

**Fig. 7.**
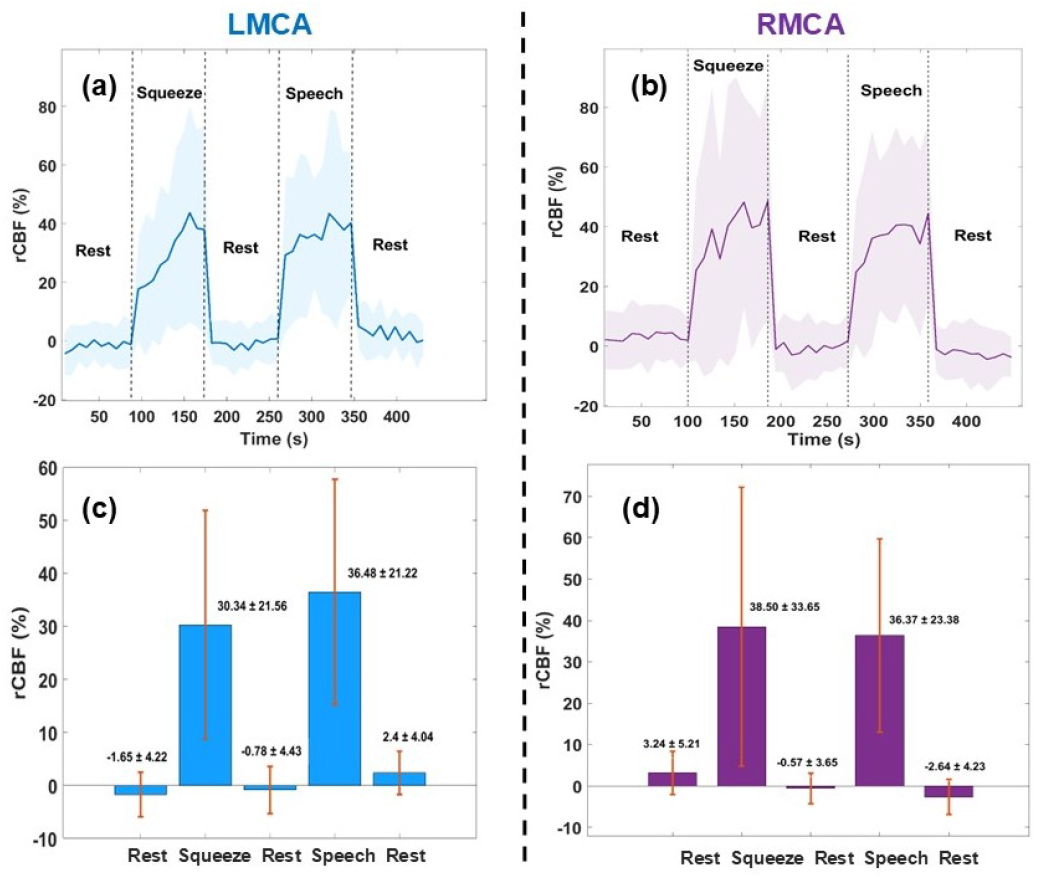
Pilot study results (n=30) for contralateral squeeze task and speech task (Q&A); (a) and (b) shows rCBF percentage, and in (c) and(d) bar graphs show mean rCBF percentage for Left and right MCA respectively.

Similarly, for RMCA, 38.50 ± 33.65% and 36.37 ± 23.38% are the measured mean rCBF values as shown in bar graphs of Fig. 7. Statistical data analysis shows a high significance (*p <* 0.001) in t-test, for squeeze task and Q&A with baseline, but no significance (*p >* 0.014) for tasks w.r.t each other as shown in Table II.

**Table II.** Significance of results for task to baseline and task to task corresponding to left and right MCA.

| Position | X1 | X2 | p value for test | Significance |
| --- | --- | --- | --- | --- |
| Left MCA | baseline | Squeeze | $4.8810^{-7}$ | Significant |
| | baseline | Speech | $6.10810^{-9}$ | Significant |
|  | Squeeze | Speech | 0.358 | Not significant |
| Right MCA | baseline | Squeeze | $1.4310^{-6}$ | Significant |
| | baseline | Speech | $6.3610^{-8}$ | Significant |
|  | Squeeze | Speech | 0.8342 | Not significant |

### D. Ischemic lacunar infarct case study

After system repeatability and accuracy evaluation in healthy volunteers, the DCS system was used at Sri Balaji Hospital, Guindy, Chennai, India for blood flow monitoring in an MCA stroke patient. The purpose of this study was to assess the suitability of the system and probe in ICU environments and to identify possible challenges in clinical translation. A 70-year-old male patient was admitted on Day 1 with complaints of difficulty in walking without support and slurred speech for one week. The initial CT scan showed bilateral MCA ischemic lacunar infarcts, as shown in Fig. 8 (a). The patient was treated with appropriate medication and discharged on Day 3. During the follow-up visit on Day 10, his condition had improved. His speech was better compared to the initial stage, although weakness was still present in the distal region of the left upper limb (UL) and left lower limb (LL). The probe was fixed on the patient’s head at temporal window (sylian fissure), as shown in Fig. 8 (b), and the experiment was carried out.

**Fig. 8.**
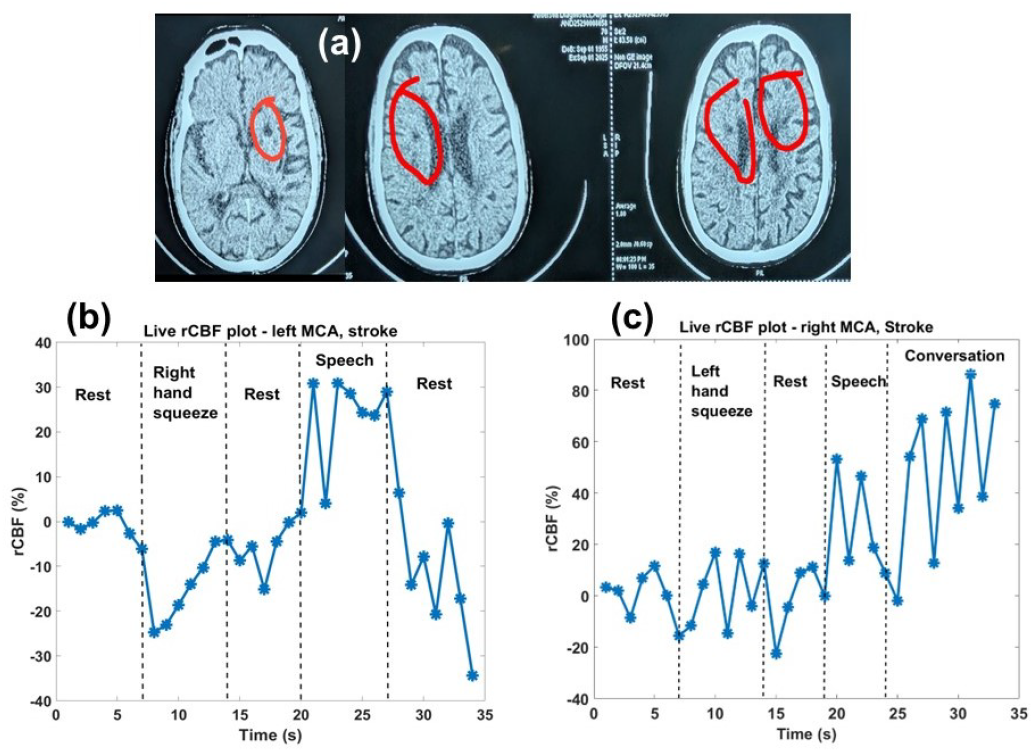
Represents (a) CT scan images showing ischemic lacunar infarcts on both hemispheres (marked with red circle), (b) and (c) shows rCBF plot for LMCA, and RMCA respectively for contralateral hand squeeze task and speech task

Fig. 8 (c) and (d) show the rCBF variation with time during contralateral hand squeeze and speech tasks for the left and right MCA, respectively. For the left MCA, the rCBF initially decreased and then increased during the right-hand squeeze task; however, the values remained close to the resting rCBF level. During the speech task, a significant change of nearly 30% in rCBF was observed. Similarly, for the right MCA, no major change was seen during the left-hand squeeze task, whereas a change of around 30% was observed during the speech task. The final segment of the rCBF plot in RMCA represents the conversation between the doctor and the patient, during which a continuous increase in blood flow was observed. These findings indicate that the developed DCS system and the probe are sufficiently sensitive for targeted and continuous blood flow monitoring in MCA and can be effectively used in different clinical and rehabilitation settings.

## V. DISCUSSION

In this study, we aimed for region-specific and simultaneous monitoring of CBF changes in MCA territories using an in-house developed DCS system with a dual probe assembly. A systematic optimal probe positioning, protocol optimization and pilot cohort study in healthy subjects has been performed. Consecutively, feasibility of the device and the method was validated in a patient affected with ischemic lacunar infract stroke.

In the optimal probe placement study, the results showed significantly larger increase in CBF corresponding to the targeted MCA region, whereas no significant change in rCBF was observed at the off-target positions (refer Fig. 5). Since, MCA traverses through Sylvian fissure, other two regions namely frontal cortical regions (Up position) and zygomatic arch (Down position), do not provide exposure window for MCA sampling. This further supports the spatial specificity of the optimized probe placement for detecting hemodynamic changes associated with the MCA territory (refer Fig. 3 (a)). Moreover, a study using Xe single-photon emission tomography [24], [25], reports a significant increase in rCBF in the contralateral motor-sensory cortex during handgrip, while no significant change was observed in the frontal region and surrounding region which supports the proposed optimal probe placement.

Furthermore, protocol optimization study was carried out to identify functional activity and speech tasks that consistently perturb CBF in the MCA. Including speech monitoring in the protocol would be helpful for assessing stroke patients with mobility impairments while their speech function remains intact. For motor activation, hand-squeeze tasks involving both ipsilateral and contralateral hand movements were evaluated. Similarly, for speech activation, different speech-related tasks such as counting, Q&A tasks were performed to determine the most suitable task capable to produce the strongest and consistent blood flow response. The motor-task results demonstrated that contralateral hand squeezing produced a significantly larger sensitivity than ipsilateral hand squeezing for both left and right MCA (refer Fig. 6 (a)). A study using TCD, reports higher flow velocity in MCA for contralateral motor activity that supports the outcomes of proposed protocol optimization study for functional activity [25].

For speech-task optimization study, the results revealed that the thinking-based speech task, i.e., Q&A task, produced the higher and consistent increase in blood flow for both left and right MCA (refer Fig. 6 (b)). This difference in CBF for counting and Q&A task is because of contribution of associated regions supplied by MCA, as the lateral frontal and parietal areas fed by the MCA assist in working memory and complex problem-solving during dialogue, and Angular Gyrus situated in the parietal lobe (supplied by outer cortical MCA branches) governs numerical processing, counting, and calculating [26]. This interpretation is consistent with a previous fNIRS study showing that high speech-planning demand produces differential activation within speech-language networks, with distinct hemodynamic responses associated with speech planning and execution [27].

Based on these observations, contralateral hand squeezing and Q&A tasks has been finalised as the motor activation and speech activation task protocol for subsequent experiments and pilot studies. The pilot study results showed a significantly higher CBF changes during contralateral hand squeezing and the Q&A speech task with respect to baseline, with consistency and reproducibility across the subjects (refer Fig. 7 and Table II). Additionally, no significant change in rCBF was observed between the motor and speech tasks. This lack of significance within tasks indicates similar blood flow demand from the motor and speech areas supplied by MCA which is also reported in [28] using near-infrared spectroscopy and transcranial magnetic stimulation. Therefore, the magnitude of CBF alone cannot be used to directly compare the two tasks or identify the task being performed. It was observed that during hand squeeze task the magnitude of CBF gradually increases and a promint change in CBF appears when subject starts feeling fatigue in hand muscles and blood supply demand in motor cortex region increases to complete the task.

In addition, we also conducted a study to determine the feasibility of using the DCS system for CBF monitoring in a clinical environment. Despite the existence of MCA ischemic lacunar infarcts, task-dependent variations in rCBF were successfully captured from both the hemispheres. Motor tasks produced minimal changes in rCBF, likely reflecting impaired functional activation associated with the stroke condition, whereas speech-related activity showed pronounced hemodynamic responses of approximately 30% in both MCAs. The gradual increase in blood flow observed during doctor–patient interaction further highlighted the sensitivity of the system to cognitive and communicative engagement. This measurement was done with single probe considering patients’ comfort on priority.

Overall, this work establishes a complete framework for MCA-specific CBF monitoring using DCS. Simultaneous bilateral MCA monitoring has the potential to provide valuable information regarding cerebral hemodynamic and recovery progression, thereby supporting clinical decision-making, specially where differences in perfusion between the affected and unaffected hemispheres may provide valuable clinical information. The future objective of this work is to further improve the temporal resolution of the system and evaluate its long-term stability, continuous bedside monitoring of cerebral perfusion in ischemic stroke patients.

## VI. Conclusion

A targeted, continuous, and simultaneous monitoring of rCBF in the MCA territories is done using in-house developed DCS system containing dual probe assembly. Optimized probe placement based on anatomical landmarks and standardized protocol enabled reliable localization of MCA-specific perfusion, while functional validation in healthy volunteers demonstrated reproducible task-evoked increases in rCBF during contralateral motor activation and speech-related cognitive tasks. Evaluation in a pilot cohort of 30 healthy participants validated the consistency and reproducibility of the system. Furthermore, preliminary monitoring in a patient with bilateral MCA ischemic lacunar infarct demonstrated the feasibility of the device for continuous neurovascular assessment in a clinics. The proposed system provides a non-invasive, targeted and operator-independent approach for real-time arterial cerebral perfusion monitoring. These findings established a strong foundation for future large-scale clinical studies and supported the potential of the DCS system for early detection of cerebral hemodynamic changes and cerebrovascular disorder in stroke patients.

## Acknowledgment

Authors gratefully acknowledge Wadhwani Research Centre of Bioengineering (Indian Institute of Technology Bombay), Technology Innovation Hub (Indian Institute of Technology Bombay), Core Research Grant (Science and Engineering Research Board), HDFC ERGO for financial support.

